# High cardiomyocyte diversity in human early prenatal heart development

**DOI:** 10.1101/2022.02.26.482029

**Authors:** Christer Sylvén, Eva Wärdell, Agneta Månsson-Broberg, Eugenio Cingolani, Konstantinos Ampatzis, Ludvig Larsson, Åsa Björklund, Stefania Giacomello

## Abstract

Cardiomyocytes play key roles during cardiogenesis, but have poorly understood features, especially in prenatal stages. Thus, we have characterized human prenatal cardiomyocytes, 6.5– 7 weeks post-conception, in detail by integrating single-cell RNA sequencing, spatial transcriptomics, and ligand–receptor interaction information. Using a computational workflow developed to dissect cell type heterogeneity, localize cell types, and explore their molecular interactions, we identified eight types of developing cardiomyocyte, more than double compared to the ones identified in the *Human Developmental Cell Atlas*. These have high variability in cell cycle activity, mitochondrial content, and connexin gene expression, and are differentially distributed in the ventricles, including outflow tract, and atria, including sinoatrial node. Moreover, cardiomyocyte ligand–receptor crosstalk is mainly with non-cardiomyocyte cell types, encompassing cardiogenesis-related pathways. Thus, early prenatal human cardiomyocytes are highly heterogeneous and develop unique location-dependent properties, with complex ligand–receptor crosstalk. Further elucidation of their developmental dynamics may give rise to new therapies.

**Graphical abstract:** 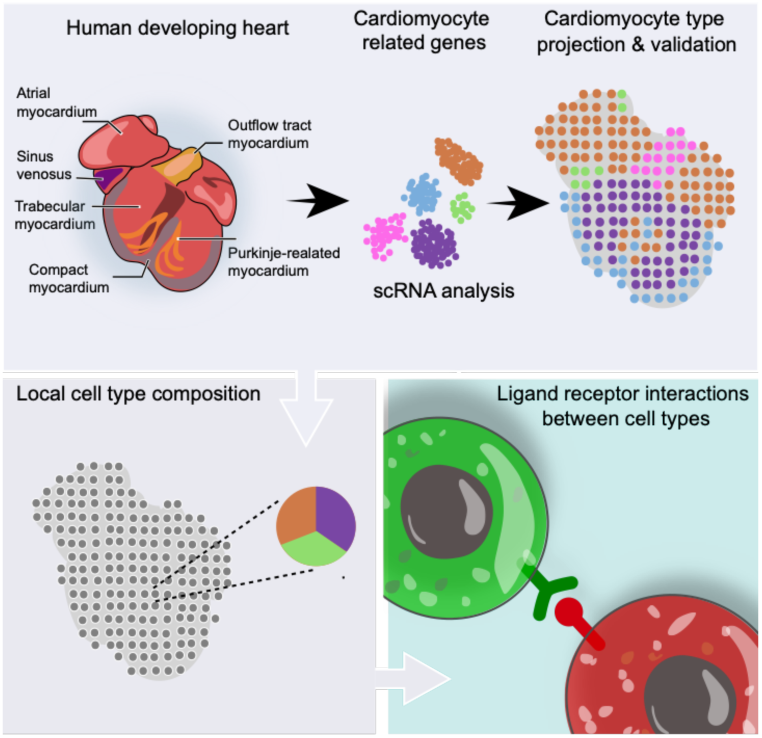

## Introduction

Cardiomyocytes (CMs) are the contracting units of the heart. Their growth and adaptation are seminal for cardiogenesis during early prenatal development, when the heart has transformed from a primitive heart tube to a growing four-chambered heart and outflow parts of the ventricles are developing (Sylva et al., 2014; Samsa et al., 2013; Sylva et al., 2014; Günthel et al., 2018). Elements of the venous pole of the cardiac tube and cardiac mesenchyme related to the sinus venosus, great cardiac veins, and pulmonary veins develop into the trabeculated atrial appendages, atrial septum, central conduit part of the atria, and sinoatrial node (SAN), from which the normal heart beat originates (DeRuiter et al., 1995; Faber et al., 2019; Soufan et al., 2004; Steding et al., 1990; Sylva et al., 2014). The atrial structures originating from the veins are myocardialized by CMs (Douglas et al., 2009; Gallego et al., 1997), which form the smooth-walled conduit parts of the atria.

However, the mechanisms involved in cell division, differentiation with development of contractility, energy production, and electrical integrity of cardiomyocytes in these developmental compartments are still unknown. Thus, exploration of CMs’ spectrum, specification, growth and interactions with other cells during early prenatal life is needed to improve understanding of cardiac health, the development of CM populations, and thus potential avenues towards new regenerative therapies for heart failure.

Asp et al. (2019) have generated the most comprehensive spatial cell atlas, at the time of writing, of human prenatal heart development between 4.5 and 9 post-conceptional weeks (PCW) using spatial transcriptomics (ST) (Salmén et al., 2018; Ståhl et al., 2016), single-cell RNA sequencing (scRNA-seq) (Zheng et al., 2017), and in situ sequencing data (Ke et al., 2013; Qian et al., 2020). This dataset forms part of the *Human Developmental Cell Atlas* (*HDCA*). At 6.5–7 PCW, Asp et al. (2019) identified three CM populations, corresponding to atrial CMs, ventricular CMs, and MYOZ2-enriched CMs. Here, we present a high-resolution characterization of the CM populations at 6.5–7 PCW, achieved by applying a computational workflow developed to dissect and characterize cell type heterogeneity, localize the cell types, and explore their ligand-receptor (L-R) interactions.

## Results

We initially applied Uniform Manifold Approximation and Projection (UMAP) (Becht et al., 2018) for dimensionality reduction and clustering of the *HDCA* 6.5–7 PCW heart scRNA-seq dataset (Asp et al., 2019) (Figure S1). To specifically investigate the CM heterogeneity of the human prenatal heart, we subclustered 726 cells corresponding to the three CM clusters identified in the *HDCA* dataset, i.e., ventricular, atrial, and *MYOZ2*-enriched CMs. Through this process eight CM clusters emerged (Figure 1A). Of these, six (designated clusters 0–2, 4, 6, and 7) were ventricular CM clusters expressing *MYH7* and two (clusters 3 and 5) were atrial CM clusters expressing *MYH6*.

**Figure 1.**
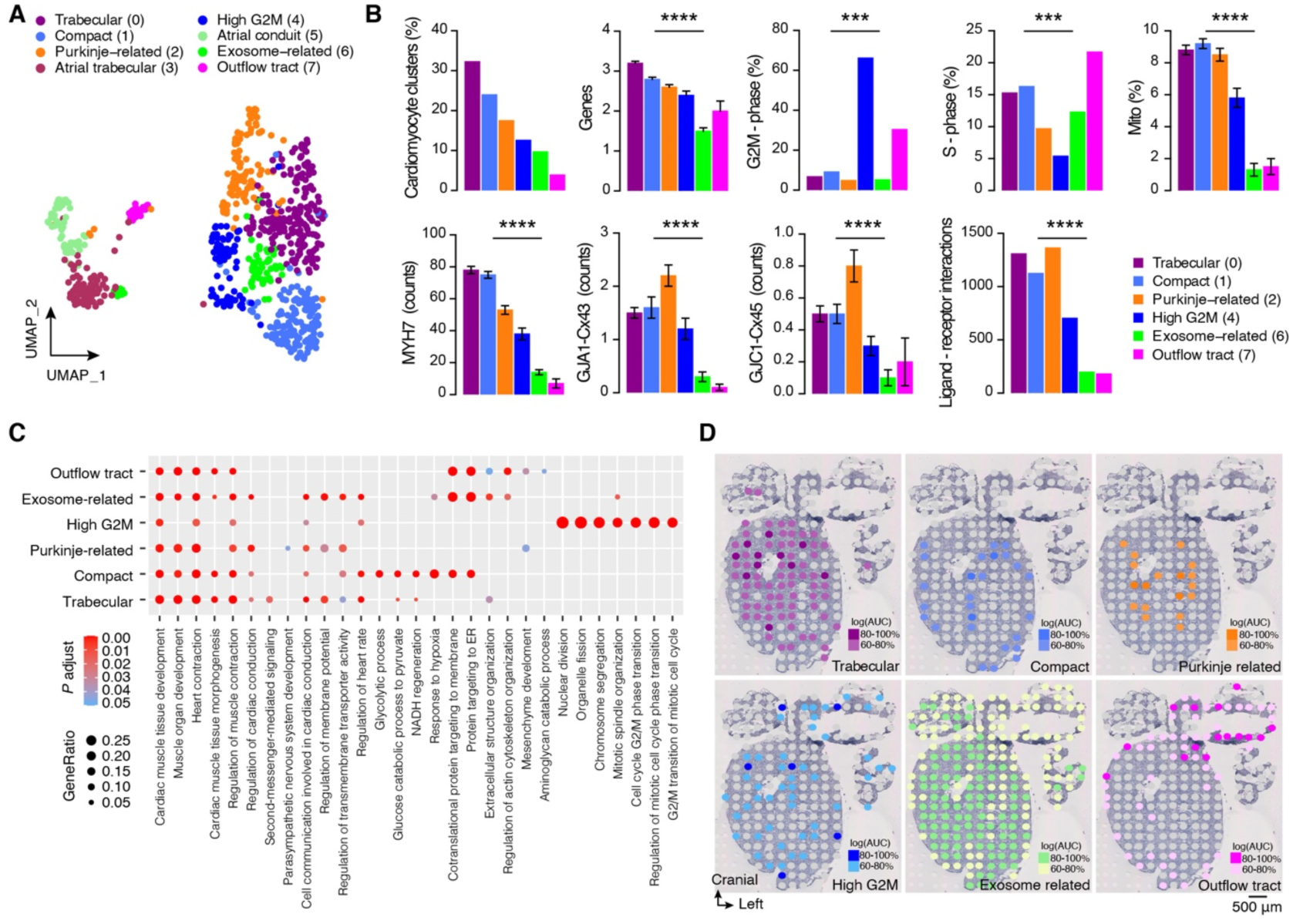
(A) Two-dimensional UMAP of scRNA-seq ventricular and atrial CM clusters. (B) Summary of basic properties of ventricular CM types: percentage of total, number of genes ×1000, percentages of cells in G2M and S phases, percentage of mitochondrial RNA, counts of *MYH7*, connexins *GJA1* (Cx43) and *GJC1* (Cx45), and number of L-R interactions. Asterisks indicate *P* values obtained from one-way ANOVA or chi-square tests on absolute numbers: *** *p* < 0.0005; \*\*\*\**p* < 0.0001. (C) GO characteristics: biological processes. (D) Deconvolution of ventricular CM types and their locations on a ST map.

### Characteristics of ventricular cardiomyocyte populations

The three largest ventricular CM clusters (clusters 0, 1, and 2; Figures 1A and 1B), contained ≈75% of the cells, with higher mitochondrial content, connexin expression and ligand–receptor (L–R) interactions than cells in the other three ventricular CM clusters (4, 6, and 7). This suggests they were more mature and morphologically integrated with higher oxidative, contractile, and electrical conductivity capacities (Kolanowski et al., 2020; Saheli et al., 2020). The third largest cluster (cluster 2) was associated with strong differential expression (DE) of the gene *IRX3*, with an adjusted *p*-value (p_val_adj) of 4.8 × 10^−17^ (Figure S2A), which is characteristic of the distal part of the Purkinje fibers (Hu et al., 2018; Kim et al., 2016). The cluster also most strongly expressed genes encoding gap junction connexins 43 and 45 (*GJA1* and *GJC1*, respectively: Figure 1B), which play key roles in fast conduction fibers (Boukens et al., 2013), and was characterized by several gene ontology (GO) terms related to conduction and signaling (Figure 1C). Taken together, these observations suggest that cluster 2 consisted of **Purkinje CMs**.

The second largest ventricular CM cluster (cluster 1) was associated with GO terms related to glycolysis (Graham and Huang, 2021) and response to hypoxia (Meng et al., 2020), indicating high energy production. Moreover, this cluster strongly differentially expressed the gene *HEY2* (p_val_adj 2.5 × 10^−27^) (Figure S2A), which is a downstream effector of NOTCH signaling that plays a pivotal role in the maturation of **compact CMs** (Firulli et al., 2020). *HEY2* represses atrial and trabecular gene expression of *TBX5* (Steimle and Moskowitz, 2017), *NPPA* and *GJA5* (encoding connexin 40). Thus, the cells in this cluster have characteristics of **compact CMs**.

The largest ventricular CM cluster (cluster 0) contained about a third of the ventricular CMs (Figure 1B). GO characteristics (Figure 1C) included more pronounced contractile- and membrane-related terms than for terms related to compact CMs. Several contractile-related genes were enriched (Figure S2A), such as *MYL2, TNNI3*, and *MYL3. HOPX*, a transcription factor involved in CM maturation (Friedman et al., 2018) was also highly expressed (p_val_adj 6.3 × 10^−26^). Thus, cells in this cluster have characteristics of **trabecular CMs**.

The smallest ventricular CM cluster (cluster 7) was associated with few L-R interactions and weak expression of both *MYH7* and connexin expression (Figure 1B). Unique GO terms were related to mesenchyme development and both extracellular and cytoskeleton traits (Figure 1C). Fractions of the cells in G2M and S phases were high: 30.4% and 21.7%, respectively.

Highly differentially expressed genes (Figure S2A) included *LGALS1* (p_val_adj 1.2 × 10^−7^), *PLAC9* (p_val_adj 5.2 × 10^−13^), *ACTG1* (p_val_adj 4.4 × 10^−12^) and *S100A11* (p_val_adj 2.9 × 10^−14^). *LGALS1* encodes a beta-galactosidase-binding protein implicated in cell–matrix interactions, and colocalizes with intracellular sarcomeric actin (Dias-Baruffi et al., 2010). *PLAC9* encodes a glycoprotein implicated in the extracellular matrix network (Cui et al., 2019). *ACTG1* encodes a major constituent protein of the contractile apparatus; and *S100A11* encodes a calcium-binding protein. These properties suggest that cells in this cluster are cardiomyofibroblasts, developing mainly in the outflow tract (**OFT CMs**), where ventricular muscle as well as semilunar valves and the great arteries develop (Boukens et al., 2009; van den Hoff et al., 2001; Sahara et al., 2019).

The second smallest ventricular CM (cluster 6) had low numbers of expressed genes and transcripts and weak expression of mitochondrial genes *MYH7* and *GJA1* (Figure 1B). Associated GO terms are indicative of high endoplasmic reticulum activities. The gene *YBX1* was highly differentially expressed (p_val_adj 8.8 × 10^−16^) (Figure S2A) and is involved in recognition of RNA molecules and exosomal transport of microRNAs (miRs) (Lin et al., 2019; Shurtleff et al., 2017; Suresh et al., 2018), in accordance with GO characteristics indicating protein targeting to ER. These **exosome-enriched CMs** may be viewed as cardiomyoblasts.

The cells in the third smallest ventricular CM cluster (**high G2M CMs)** had fewer expressed genes and transcripts, and expressed fewer mitochondrial genes than compact or trabecular CMs (Figure 1B). GO terms (Figure 1C), and high differential expression of such genes as *PTTG1* (p_val_adj 3.9 × 10^−35^) and *TUBA1B* (p_val_adj 9.7 × 10^−29^), indicated pronounced G2M cell cycle activity. These **high G2M cardiomyoblasts** may be viewed as an intermediate stage between exosome-enriched CMs and compact, trabecular, and Purkinje CMs.

Gene expression and cell cycle phase scores were highly variable both between and within the ventricular CM clusters (Figure S2B, C). All ventricular cell types included some in the G2M phase. Mitochondrial gene expression was also highly variable. It was low in OFT CMs, but there were also distinct fractions of cells with low expression in compact, trabecular, and exosome-enriched CMs (Figure S2B). These weakly-expressing cells were localized in the HDCA dataset to the *MYOZ2*-enriched CM cluster, while the remaining cells were localized to the ventricular CM cluster. Overall, the weakly-expressing cells (16% of the total) had lower numbers of expressed genes and transcripts and weaker mitochondrial gene expression than other cells, but a higher fraction in the G2M phase (26% vs 14%; *p* < 0.001 according to the chi-square test). These findings suggest that these cells might be cardiomyoblast-like.

As additional validation, the CM scRNA-seq clusters were deconvoluted onto ST cardiac morphology maps, where four of the six ventricular CM clusters were anchored to relevant anatomic structures of the heart (Figure 1D and Figure S3). Compact CMs were mainly located close to the epicardium of both ventricles and in the septum, while trabecular CMs were located in the inner parts of both ventricles. Purkinje fiber-related CMs were localized to the distal-central part of the right and left ventricles, while OFT CMs were mainly located in the outflow tract. Exosome-related and high-G2M CMs were diffusely distributed in the ventricles.

### Atrial cardiomyocyte types

As there were too few **SAN CMs** to identify by the clustering algorithm, they were detected using a supervised approach as cells coexpressing *TBX18, SHOX2*, and *HCN4* (Easterling et al., 2021; Liang et al., 2017; Wiese et al., 2009). In the map generated by UMAP analysis (Figure 2A), showing the two atrial clusters (clusters 3 and 5), SAN CMs are located at the upper end of atrial cluster 5. Figure 2A also shows the expression sites of 10 markers related to atrial function. *TBX18* and *SHOX2* were only expressed in cluster 5: *TBX18* specifically at the upper end in the SAN_CM area, while *SHOX2* expression was more widely dispersed within cluster 5. *SHOX2* was prominently expressed in both the SAN (p_val_adj 2.1 × 10^−23^) and cluster 5 (p_val_adj 1.2 × 10^−69^) (Figure S4). At this developmental stage expression of *SHOX2*, a growth factor of mesodermal origin, is restricted to the right atrial conduit area (Espinoza-Lewis et al., 2009; Hu et al., 2018; Liu et al., 2012) and left atrial conduit area around the pulmonary vein orifices. Our results confirm that, in concert with the proepicardial and mesenchymal derived transcription factor *TBX18* (Cai et al., 2008), *SHOX2* participates in development of SAN pacemaker CMs through suppression of the myocardial contractile gene program’s expression. This involves suppression of the early cardiac transcription factor *NKX2-5* (expressed less strongly in cluster 5 than cluster 3, and in SAN less strongly than in cluster 5 CMs: *p* < 0.008 and < 0.003, respectively), resulting in upregulation (*p* < 0.0001) of conduction-associated transcription factor *TBX3*. Consequences include downregulation of *NPPA* (expressed less strongly in cluster 5 than in cluster 3 and less strongly in SAN than in cluster 5 CMs: *p* < 0.0001 and < 0.004, respectively) and the connexin 40-encoding gene *GJA5* (expressed less strongly in cluster 5 than in cluster 3: *p* < 0.0001). This prevents SAN CMs from becoming contracting fast-conducting atrial CMs (van Eif et al., 2019; Wiese et al., 2009). Expression of *PPP1R1A* was higher (*p* < 0.0001) in the SAN than in cluster 5. This gene encodes a cytoplasmic protein phosphatase inhibitor involved in signal transduction and carbohydrate metabolism.

**Figure 2.**
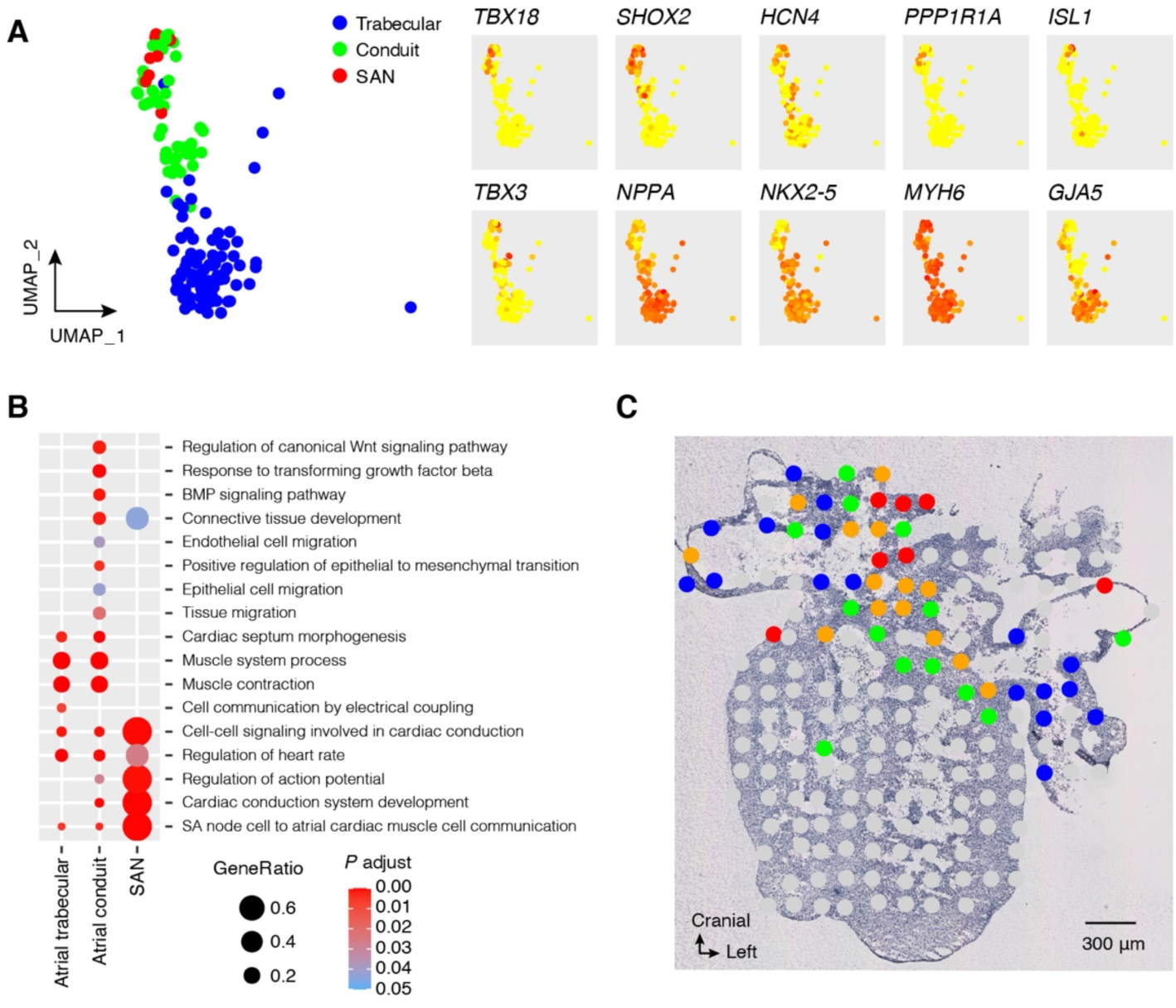
Atrial CMs. (A) Two-dimensional UMAP of atrial trabecular, conduit, and SAN CMs and heat maps of relevant genes. (B) GO characteristics: biological processes. (C) Deconvolution of atrial CM types and their locations on a ST map.

The two atrial CM clusters (clusters 3 and 5) were similar in muscular contractility GO terms (Figure 2B), but only the smaller cluster (5) expressed epithelial cell migration, epithelial to mesenchymal transition, endothelial cell proliferation, BMP, and canonical WNT signaling pathways. These differences suggest that the larger cluster (3) is related to the primordial atria, developing mainly into the trabeculated atrial appendages (**atrial trabecular CMs**) while cluster 5 is related to the central smooth-walled part of the atria developing from the epicardium and mesenchyme and large veins, including the pulmonary veins (Douglas et al., 2009; Gallego et al., 1997; van den Hoff et al., 2001) (**atrial conduit CMs**). The GO terms of SAN CMs (Figure 2B) were related to conduction, regulation of action potential, and heart rate. As additional validation, Figure 2C and Figure S3 show that SAN CMs were localized to the upper part of the right atrium and the conduit atrial CMs mainly localized to the central part of the atria, while the trabecular atrial CMs were located more peripherally in the atria. Signals of both trabecular and conduit CMs were detected in several central spots. This was presumably because each spot in the spatial grid of a microdissected transcriptomic map may represent 20– 30 cells, which could, of course, include different colocalized cell types.

### Spatial colocalization of different cell types

The interactions between different cell types during organ development are highly important. By deconvoluting non-cardiomyocyte (non-CM) single-cell clusters from the *HDCA* (Figure S1) onto ST spots, we can visualize colocalized cell types and thus obtain indications of the key cells and their interactions in local development. Figure 3A shows the ST spots in the OFT region of one tissue section that contained OFT CMs and Figure 3B the fraction of spots with non-CMs colocalized to these OFT CM spots. Fractions of epicardium-derived cells, fibroblast-like cells related to larger and smaller vasculature, fibroblast-like smooth muscle cells and cardiac skeleton cells, endothelial and Schwann progenitor cells were all high in these spots. In ST spots with unique atrial conduit CMs (Figure 3B) a similar composition of cell types was observed although also epicardial cells were present. Fractions of non cardiomyocyte cell types were lower. This indicates that CMs may play a less prominent role in the OFT region (where major developmental events include septation and formation of the aorta, pulmonary artery and semilunar valves) than they play in the atria, where muscle formation is the major developmental event.

**Figure 3.**
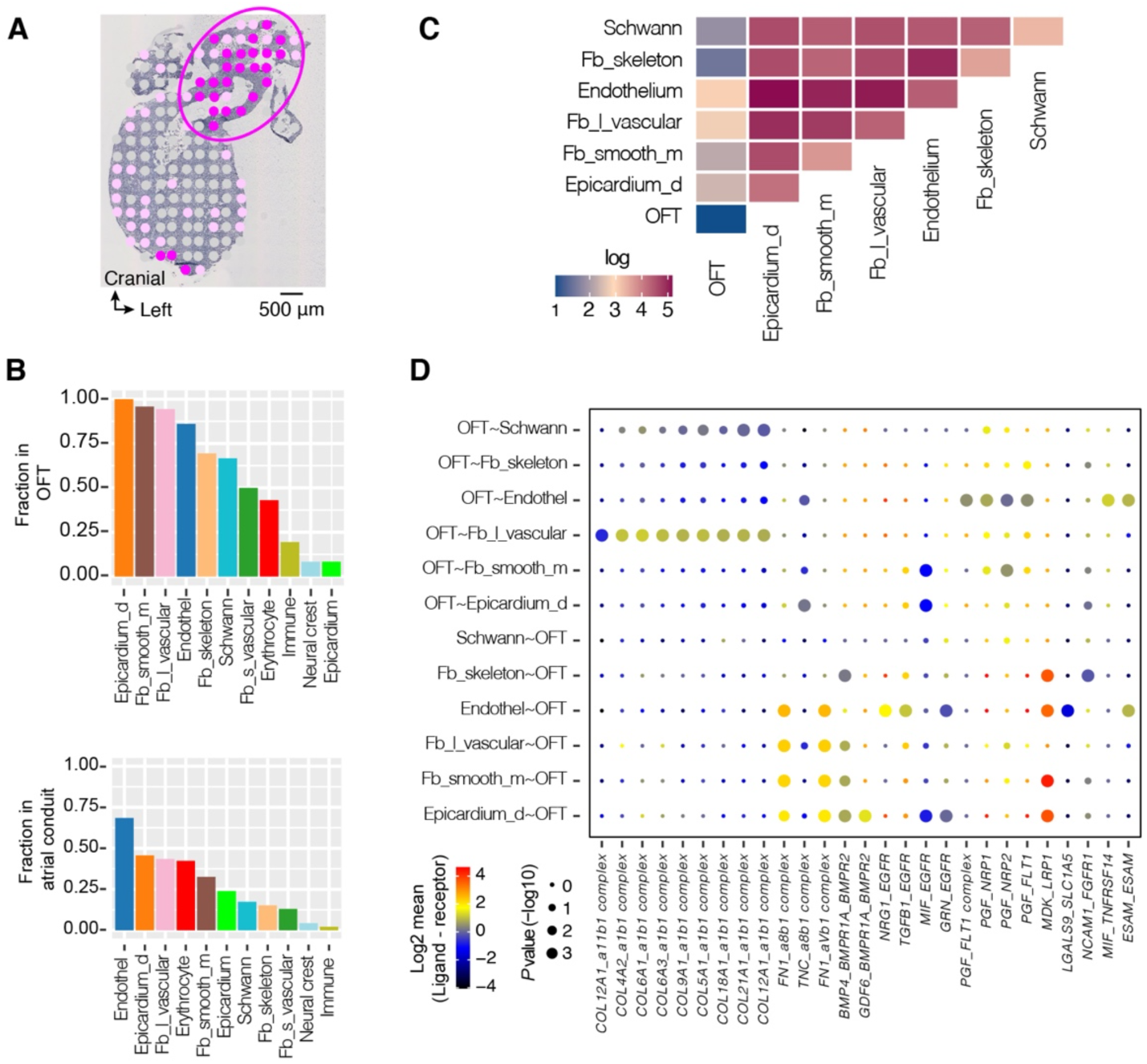
(A) ST spots with projected OFT CMs in the OFT region (ellipse) in heart section 115. (B) Fraction of spots expressing non-CM cell types deconvoluted from scRNA-seq clusters described in Figure S1 for spots expressing (above) OFT CMs and (below) conduit atrial CMs. (C) Heat map showing numbers of L-R interactions between OFT CMs and non-CMs. (D) Specific highly expressed L-R crosstalk between OFT CMs and colocalized non-CM cell types. Log2mean indicates the L-R ratio. Abbreviations of non-cardiomyocyte cell types: Epicardium-d: Epicardium-derived, Fb_l_vascular: Fibroblast-like (larger vascular development), Fb_smooth_m: Fibroblast-like smooth muscle cells, Schwann: Schwann progenitor cells, Fb_skeleton: Fibroblast-like (cardiac skeleton).

### Ligand–receptor crosstalk

The number of L-R interactions (Efremova et al., 2020) between different CM types was low, with practically no integrin interactions (Figure S5A). By contrast, interactions between ventricular CM and non-CM cells (Figure S5A) were pronounced and highest for the regionally located compact, trabecular, and Purkinje-related CMs, which also had the highest number of integrin interactions, indicating the importance of connective tissue and vascular structures in cardiac development and in CM development and anchoring.

In OFT CM spots, the majority of detected L-R interactions (Figure 3C) were between fibroblast-like cells related to vascular development, endothelial cells, epicardium-derived cells and Schwann progenitor cells. Crosstalk between OFT CMs and these cell types was comparatively weak, possibly because CMs play a minor role in OFT development, which involves septation and valve formation. Of 27 OFT CM interactions (Figure 3D) identified, 12 involved integrins with specific L-R interactions from and toward the OFT CMs. Of non-integrins, OFT CMs expressed placental growth factor, which acts at several receptors (FLT1, NRP1, NRP2) stimulating endothelial growth and angiogenesis. More non-CM ligands acted on OFT CMs than vice versa.

The complexity of L-R interactions is examplified by those involving epidermal growth factor receptor (EGFR), which participates in crosstalk between various non-CM cell types and OFT CMs depending on the ligand involved (NRG1, TGFB1, MIF, or GRN). This is consistent with EGFR’s role in the development of bicuspid valves and vascular structures (Makki et al., 2013; Schroeder et al., 2003). Other agents involved in the interactions included BMP, LRP1 (low density lipoprotein receptor) and TNFR. BMP is a member of the TGF-beta superfamily and, along with NOTCH signaling, regulates CM proliferation (D’Amato et al., 2016; Sorensen and van Berlo, 2020). LRP1 affects cardiomyocyte lipid uptake, while TNFR activation may be involved in apoptosis, and has been described as recruiting cardiac progenitor cells (Allukian et al., 2013) and stimulating OFT CMs. LGAL, NCAM1 and ESAM are ligands involved in cell to cell and cell to matrix interactions. Thus, the identified LR crosstalk reveals fundamental aspects of the development of OFT CMs in their environment.

The exosome-enriched CMs were interspersed throughout the ventricular muscle (Figure 1D). Only 32 detected L-R interactions of these cells were significant, of which 11 involved integrins (Figure S5B). Integrin ligands were only expressed at exosome-enriched CMs, indicating unidirectional binding to non-CMs. Both FGF and VEGF ligands acted through various receptors at colocalized cells, as did TNF, BMP, NPPA, and thyrosine kinase AXL, affecting the smooth muscle, fibroblasts, and endothelium. Extracellular instructive signals affecting exosome-enriched CMs included L-R crosstalk with the nodal morphogen TGF-beta pair LEFTY2_TDGF1 (Barnes and Black, 2016; Behrens et al., 2012; Noseda et al., 2011) and CD47_SIRPA, suggesting that these CMs are at an early stage of development. SIPRA is a cell surface receptor expressed at cardiomyocyte progenitors and early cardiomyocytes (Dubois et al., 2011; Skelton et al., 2014).

### Sinoatrial node and conduit cardiomyocytes

Figure 3B shows that ST spots with atrial conduit CMs contained, with the addition of epicardial cells, similar non-CM cell types to spots with OFT CMs, although with smaller non-CM fractions, suggesting that the major cell type in these spots was conduit CMs. There were only five ST spots with SAN CMs, which prevented meaningful analysis of their cell composition. As the SAN is part of the conduit atrium, it is thought that the composition of SAN spots is similar to that of conduit CM spots. The composition of cell types is similar to that observed in studies of dissected embryonic mouse atrium containing the SAN (Goodyer et al., 2019; Li et al., 2019).

We detected more L-R interactions of conduit and SAN CMs with non-CM cells (431, including 138 integrin interactions) than those of OFT CMs. Our data show that collagen L-R pairs communicate via CM-type specific isoforms. Conduit CMs communicated uniquely with NPPA with endothelium, epicardium, smooth muscle and Schwann progenitor cells (Figure 4B) and SAN CMs communicated uniquely via PDGFB. Several interactions were specific for SAN, including those involving NOTCH, EGFR, TGFB, and WNT members that play essential roles in cardiogenesis (MacGrogan et al., 2018). The only WNT family member expressed in this context was WNT5, which primarily acts through the non-canonical WNT pathway (Liang et al., 2020; Pahnke et al., 2016), although it may also play a role in the canonical pathway (Grumolato et al., 2010).

**Figure 4.**
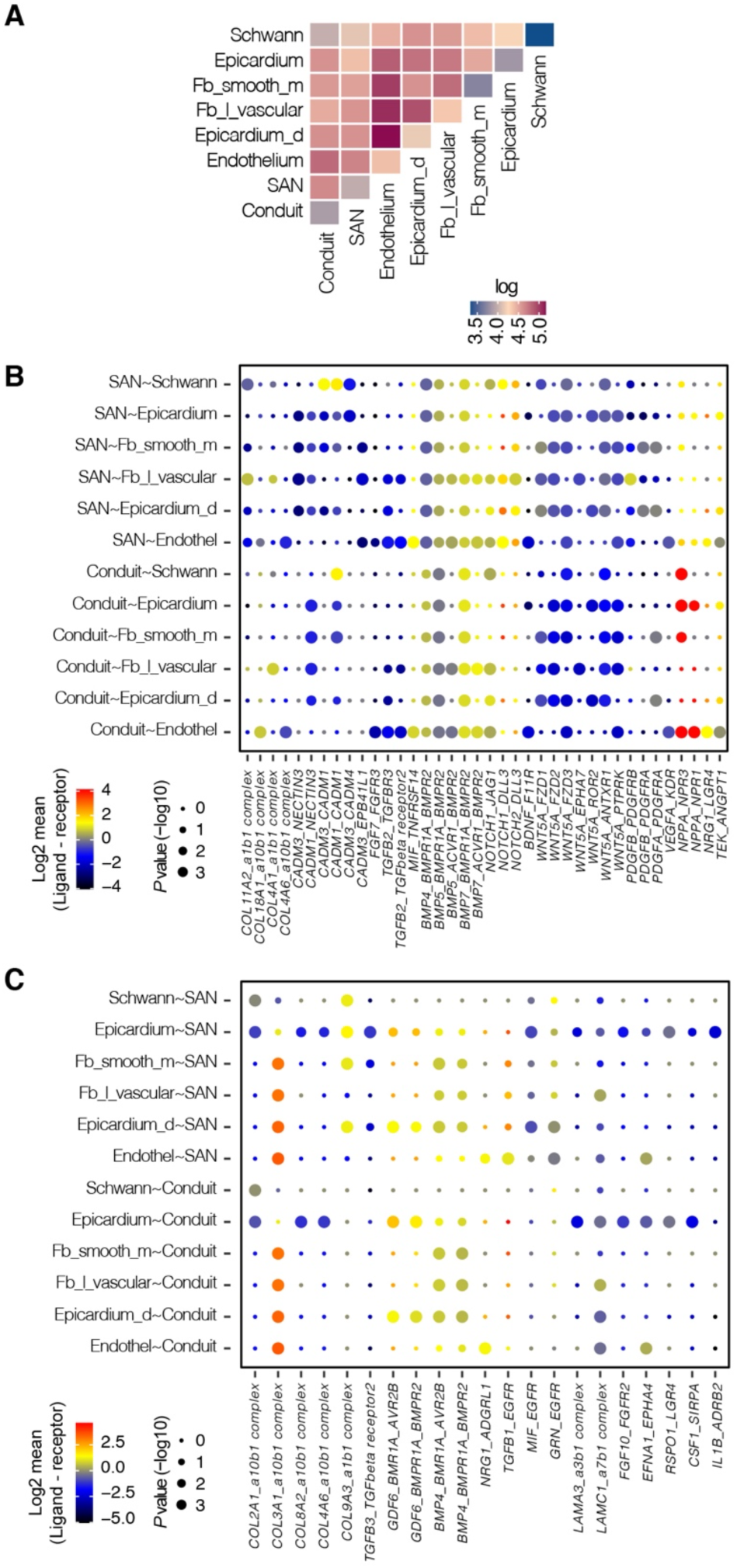
(A) Heatmap showing numbers of L-R interactions between conduit and SAN atrial CM clusters and six non-CM clusters. (C) Specific highly expressed L-R crosstalk between conduit and SAN CMs and colocalized non-CMs. (D) Specific highly expressed L-R crosstalk between colocalized non-CMs and conduit and SAN CMs. Log2mean indicates the L-R ratio. Abbreviations of non-cardiomyocyte cell types: Fb_l_vascular: Fibroblast-like (larger vascular development), Epicardium-d: Epicardium-derived, Fb_skeleton: Fibroblast-like (cardiac skeleton). Fb_smooth_m: Fibroblast-like smooth muscle cells, Fb_s_vascular: Fibroblast-like (smaller vascular development).

TGFB and BMP2 are reportedly involved in formation of the SAN (Easterling et al., 2021). We found that several TGFB L-R interactions affect SAN CM specifically, as well as BMP2 and BMP4. In addition, both conduit and SAN CMs communicate with their environment via BMP5 and BMP7, while BMP4, expressed in epicardium-derived cells, smooth muscle, and fibroblasts, acts on these CMs. Epicardial cells that have broad L-R interaction with non-CM cells, especially epicardium-derived cells (Figure 3B), also interact uniquely with both conduit and SAN CMs via specific integrins L-R pairs and LAMA3, FGF1, EFNA1 and LGR4 L-R pairs (Figure 3D). These ligands are instrumental in epithelial-to-mesenchymal transition (Ardila et al., 2021; Huang and Chen, 2021; Itoh, 2016; Zhang et al., 2020) that may in the atrial wall of venous origin contribute to its myocardialization (Cai et al., 2008; Greulich et al., 2011; Singh and Epstein, 2013; Zhou et al., 2008). Interestingly epicardial cells also uniquely interact with SAN cells with interleukin 1b acting at the adrenoreceptor B2.

## Discussion

The human embryonic heart at 6.5–7 PCW is in a dynamic phase, in which the left and right ventricles of the myocardium grow and the musculature of the proximal OFT is still developing. Our results show that rates of cell division and RNA synthesis are high, but heterogeneous, in the main part of the ventricular CM population, and they have high heterogeneity in expression of mitochondrial genes and connexin 43 (a marker of the intercellular electrical connectedness between CMs). This heterogeneity is consistent with previous findings that cell cycle gene expression downregulates sarcomeric and cytoskeletal markers, through location-specific signaling molecules that may influence the proliferation of colocalized cells (Li et al., 2019). We also detected extensive L-R crosstalk, with specific profiles, as exemplified for OFT, exosome-enriched, conduit atrium, and SAN CMs.

Two cardiomyoblast-like cell types are dispersed throughout the spatial compartments. The more immature, small exosome-enriched CMs have traits of an immature CM type that can develop into more differentiated CM types. These CMs may warrant more attention in the development of exosome-based regeneration of CMs for therapeutic heart failure treatments (Barile et al., 2017). High G2M CMs, the other type of cardiomyoblast-like cells, have gene expression characteristics more like clearly located CM types, including strong expression of *MYH7* and only slightly weaker expression of mitochondrial genes and *GJA1*. Thus, these CMs have traits of more mature CMs than the exosome-enriched CMs.

The OFT CMs differ from the compact and trabecular CMs, having a more fibroblast-like profile. The proximal OFT, conus of the right ventricle and left ventricular OFT develop later than the compact and ventricular myocardium (van den Hoff and Wessels, 2020). With respect to DE and GO characteristics, the OFT CMs expressed genes related to connective tissue development. There is some debate as to whether outflow myocardialization is based on the ingrowth of ventricular CMs or differentiation from mesenchyme, epicardium, and endocardium related to the OFT (van den Hoff and Wessels, 2020). Our results suggest that the latter source may be important, at least in the hearts analyzed.

In conclusion, our combined analysis of single-cell and spatial transcriptomic analysis, as well as L-R crosstalk between spatially colocalized cell types, reveals a complex landscape of dynamic changes in the cellular composition, expression patterns, and functions of CMs in the early prenatal human heart. Our approach also provides a strategy to characterize further the combined and publicly available single-cell and spatial transcriptomics resources.

## Acknowledgment

This work was supported by the Swedish Research Council as Formas grant 2017-01066 and Vetenskapsrådet grant 2020-04864 to S.G., the Erling-Persson family foundation and the Knut and Alice Wallenberg Foundation. A.B. was financially supported by the Knut and Wallenberg Foundation as part of the National Bioinformatics Infrastructure Sweden at SciLifeLab.

## Author contributions

C.S. conceived and designed the study, analysed the data, interpreted the results, wrote the manuscript and designed the figures. E.W. performed tissue sectioning and guided histological annotations. A.M.B. assisted in biological interpretation. E.C. conceived the study and assisted in biological interpretation. K.A. created the artwork. LL analysed the data and interpreted the results that contributed to the final manuscript. Å.B. assisted in bioinformatics analysis. S.G. conceived the study, guided data analysis, interpreted the results, wrote the manuscript and designed the figures. All authors helped with manuscript preparation.

## Supplemental information titles and legends

Figure S1. UMAP analysis on previously published single cell data derived from a human fetal heart at 6.5 PCW.

Figure S2. A: Top 12 differentially expressed genes in the ventricular CM clusters. B: Characterization of ventricular CM types as a function of percentage of mitochondrial RNA. Upper panel: number of transcripts, G2M.Score, and S.Score. Lower panel: *MYH7* and *GJA1* (connexin Cx43) transcripts in the six ventricular CM clusters. Origins of cells from base clusters are indicated: (1) ventricular CM; (13) *MYOZ2*-enriched CM. C: The origin of CMs in base cardiomyocyte clusters (ventricular, atrial, and *MYOZ2* CMs) and their fractions in cell cycle G1, G2M, and S phases.

Figure S3. scRNA-seq CM clusters projected on the nine hematoxylin-eosin image ST maps.

Figure S4. Top 18 differentially expressed genes in the atrial CM clusters and SAN cells.

Figures S5. A: Number of L-R interactions between ventricular CMs and ventricular CMs (top panel) and non-CM (bottom panel) scRNA-seq clusters described in Figure S1. Yellow and blue indicate non-integrin and integrin interactions, respectively.

B: L-R crosstalk between exosome-enriched CM and non-CM cell types. Log2mean indicates the L-R ratio.

Figure S6. Deconvolution of scRNA-seq determined clusters on ST spots.

Comparison of methods: Stereoscope (Andersson et al., 2020) and AUC calculated cluster profiles based on the 12 top differentially expressed genes for each cluster. (A) AUC_15_ correlated to Stereoscope_15_. (B) Stereoscope_15_ correlated to Stereoscope_15_. (C) AUC_15_ correlated to AUC_15_. (D) AUC_17_ correlated to AUC_17_. (E) AUC-CMs correlated to AUC-CMs. Data shown in (A) to (C) are based on 15 tsne-determined scRNA-seq clusters (Seurat 2.3.4) (Asp et al., 2019). Data shown in (D) are based on 17 UMAP-determined scRNA-seq clusters (Seurat 4.0.0) used in this work. (E) Crossed correlations show correlation with *p* ≥ 0.01.

## STAR*METHODS

### KEY RESOURCES TABLE

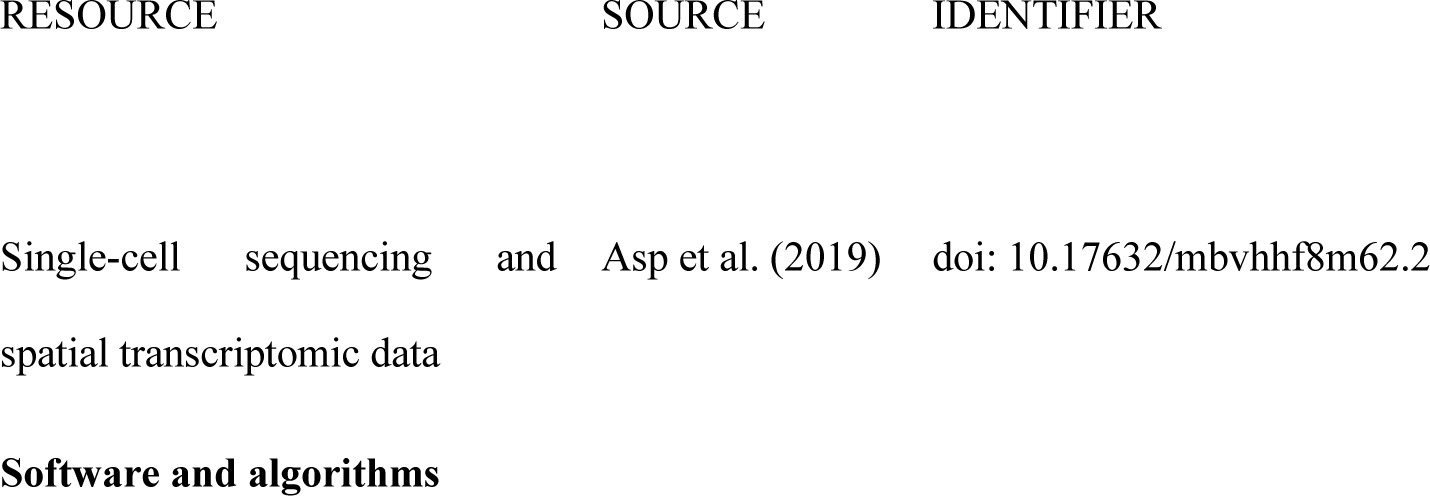
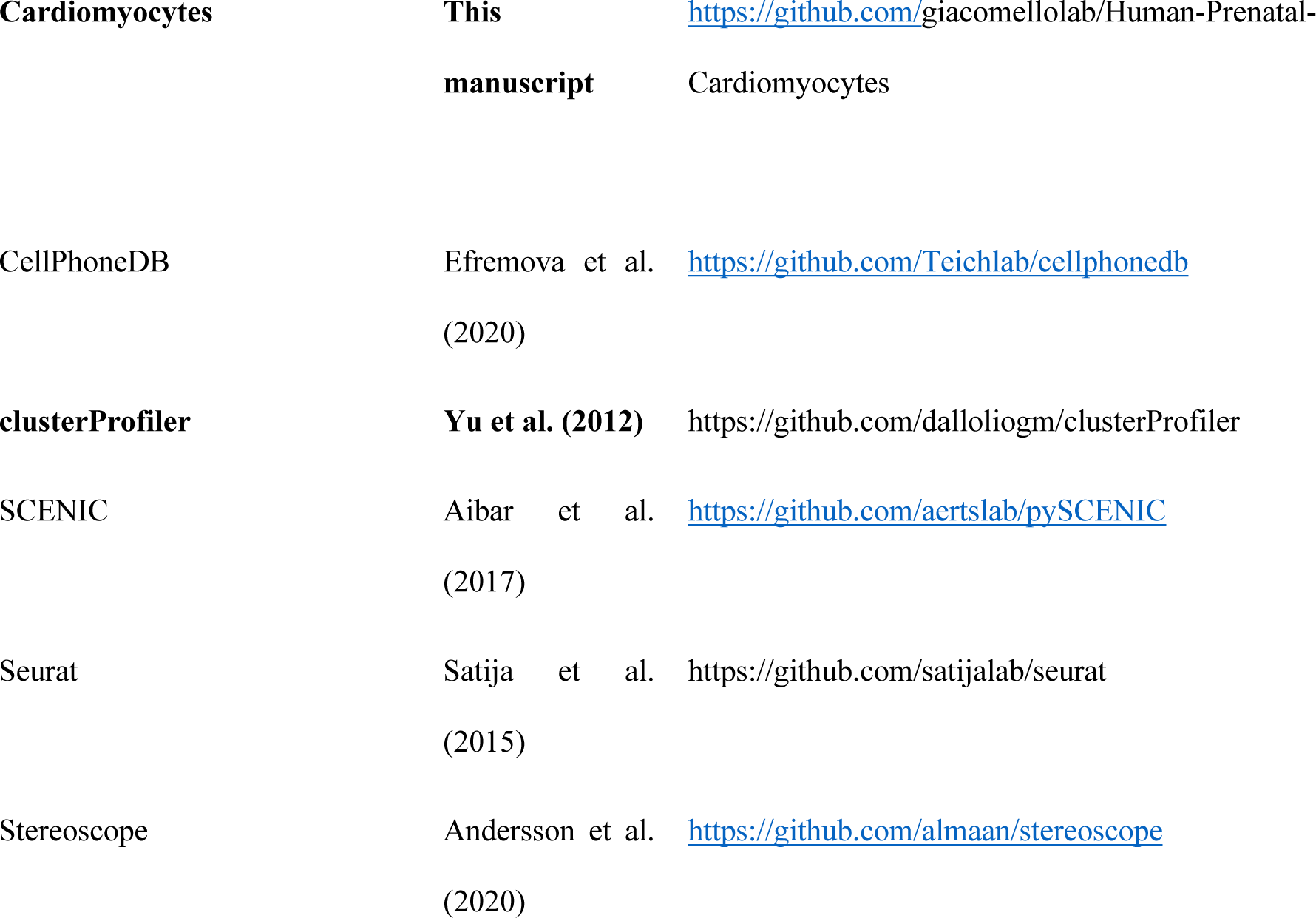

### RESOURCE AVAILABILITY

#### Lead contact

Requests for further information and/or resources and reagents should be directed to and will be addressed by the lead contacts: Dr. Christer Sylvén and Dr. Stefania Giacomello.

#### Materials availability

This study did not generate new materials.

#### Data and code availability

- scRNA seq and spatial transcriptomics data are available at Asp et a. (2019) and doi:10.17632/mbvhhf8m62.2. Background ST maps grobs data are available at https://giacomellolabst.shinyapps.io/hpc-shiny/
- Code is available at https://github.com/giacomellolab/Human-Prenatal-Cardiomyocytes
- Any additional information required to reanalyze the data reported in this paper is available from the lead contact upon request.

### METHOD DETAILS

#### Data

The exploration of prenatal CM phenotypes was based on sc-RNAseq and ST data from the *HDCA* (Asp et al., 2019). In summary, data obtained from analyses of two hearts were analyzed. Data relating to one (of 6.5–7.0 PCW) were used for single-cell transcriptomic (scRNA-seq) analysis and data relating to the other (of 6.5 PCW) for ST analysis. These two hearts were considered to be biological replicates.

Using a 10X Genomics Chromium workstation, 3717 single-cell transcriptional profiles, with on average transcripts of 2900 genes and 11,000 unique transcripts per cell, were generated after quality trimming and filtering. The ST dataset consisted of 1515 individual spots (i.e., data points equivalent to microdissections, each containing ≈ 30 cells, with on average transcripts of ≈2100 genes and ≈4800 transcripts per cell.

The heart was sectioned consecutively in the transversal plane and ST analysis was performed on nine tissue sections at levels 0, 1, 115, 200, 360, 361, 520, 521, and 675.

#### scRNA-seq analysis

The scRNA-seq analysis was performed using the Seurat package, version 4.0.0 (Butler et al., 2018). The whole material was analyzed first, then the three cardiomyocyte clusters were subclustered under the same conditions. A subset of cells expressing between 200 and 6000 features, with <50,000 counts and <20% mitochondrial content, was produced. The SCTransform function was used to regress out the influence of percent.mt, S.Score, and G2M.Score. Thereafter, 3675 cells remained for further analysis. After PCA, the FindNeighbours (dims = 1:20) and FindClusters (resolution = 0.40) routines were run. For cluster visualization, we used UMAP (UMAP) (dims = 1:20). The three muscle clusters were subset and the Seurat analysis was rerun as before. The FindClusters routine was run, with 0.8 resolution. SAN cells were identified by the simultaneous expression of *SHOX2, TBX10*, and *HCN4* (Easterling et al., 2021; Liang et al., 2017; Wiese et al., 2009). To identify differentially expressed genes, pairwise comparisons of individual clusters against all other clusters were performed using the FindAllMarkers routine (settings: min.pct = 0.25; logfc.threshold = 0.25).

#### GO characterization

GO characteristics of gene clusters were determined using the clusterProfiler package (version 3.14.3) (Yu et al., 2012) for all DE genes with an average value of logFC above zero, and adjusted *p* <0.01. The compareCluster function was used, with pvalueCutoff = 0.05.

#### Cardiomyocyte cell type validation

The cardiomyocyte types characterized by GO and differential gene expression profiles were validated according to their spatial expression in spatial microdissected transcriptomic maps with barcoded regions of 100 µm in diameter that limited the resolution to ≈10–40 cells. The grid was projected on the tissue section image, enabling morphological localization of the cluster expressions.

To deconvolute spatial locations of the CM clusters, identified using scRNA-seq data, and the original non-CM single-cell clusters from the *HDCA* in the ST map, we used the AUCell algorithm, which identifies enriched gene sets in scRNA-seq data (Aibar et al., 2017). For each CM cluster identified using scRNA-seq data, we defined the active gene set as the top 12 differentially expressed genes or the marker genes for the SAN. The **AUCellbuildRankings** function was applied to the ST dataset, to build the “rankings” for each ST spot, i.e., expression-based ranking for all the expressed genes in each spot. Then the **AUCellcalc AUC** function was used to calculate whether a critical subset of the input gene set was enriched within the expressed genes for each ST spot. The AUC function thus represents proportions of expressed genes in the ST spot, and their expression relative to other genes in it. In this way, the population of cells present in the ST spot can be explored, according to the gene set’s expression. To validate the method, cross-correlation was applied to the same scRNA-seq and ST data analyzed using the Stereoscope method (15 tsne-determined clusters with Seurat 2.3.4) (Andersson et al., 2020; Asp et al., 2019). Figure S6 shows correlations between Stereoscope and AUC (A), Stereoscope and Stereoscope (B) and AUC and AUC (C) data. Figure S6D shows AUC-AUC correlation for the 17 clusters identified with UMAP (Seurat 4.0.0) in this work. Figure S6E shows AUC-AUC correlation for the nine muscle clusters analyzed in this study. Crosses indicate correlations with *p* ≥ 0.01. Figure S6 shows that deconvolution using the AUC method discriminates between cells in a manner similar to the Stereoscope method and that the different CM types can be deconvoluted.

As dynamics of ventricular and atrial CMs differ, results for the two types of CM are presented separately.

#### L-R crosstalk

The L-R crosstalk between different cell types was analyzed using the cellphoneDB package (Efremova et al., 2020). The standard method was used. Result precision was set to 2. After 100 iterations, L-R pairs between cell types were identified, with significance set at *p* < 0.01.

#### Quantification and statistical analysis

Analysis was done with the R Stats package. Continuous data were analysed with Student’s t test or ANOVA followed by pairwise.t test. Categorical data were analysed with ChiSquare test or Kruskal Wallis test.

#### Code availability

All scripts written for the analyses presented in this paper are available at the following Github link: https://github.com/giacomellolab/Human-Prenatal-Cardiomyocytes

