## Supplementary figures and images for "High cardiomyocyte diversity in human early prenatal heart development"

### Supplementary figures

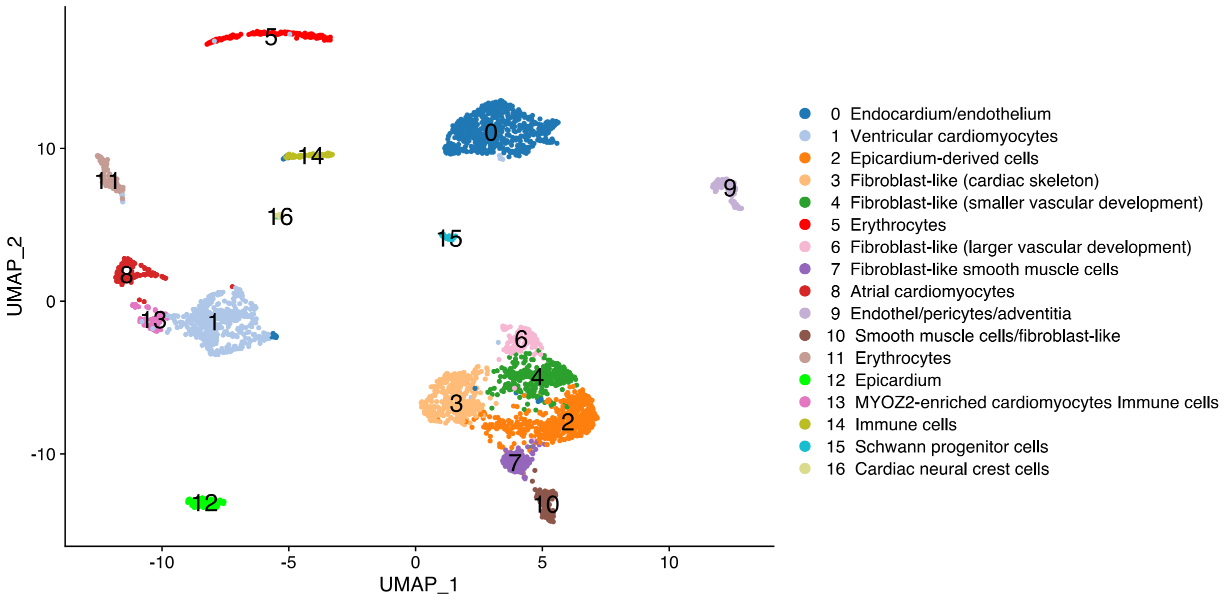

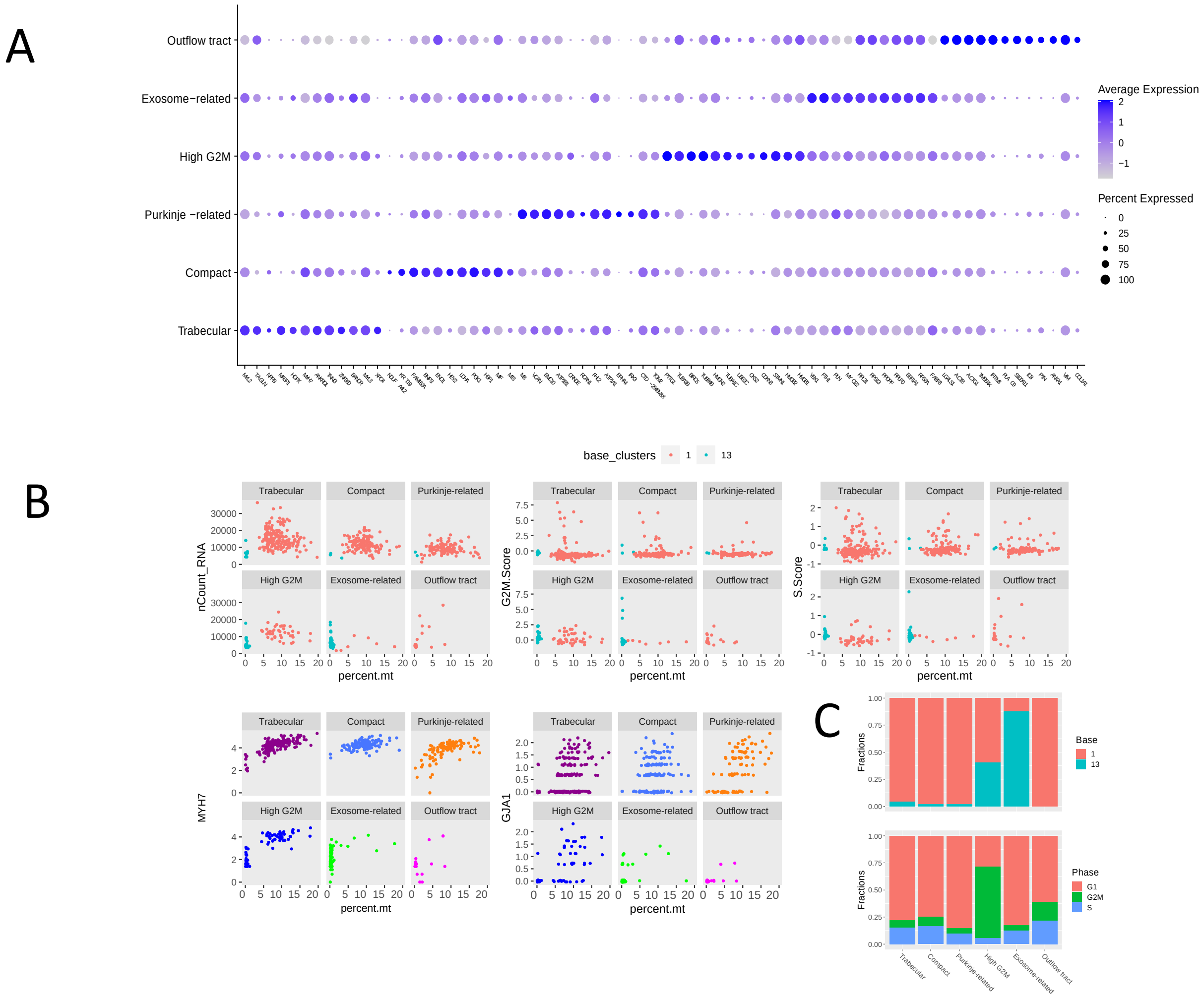

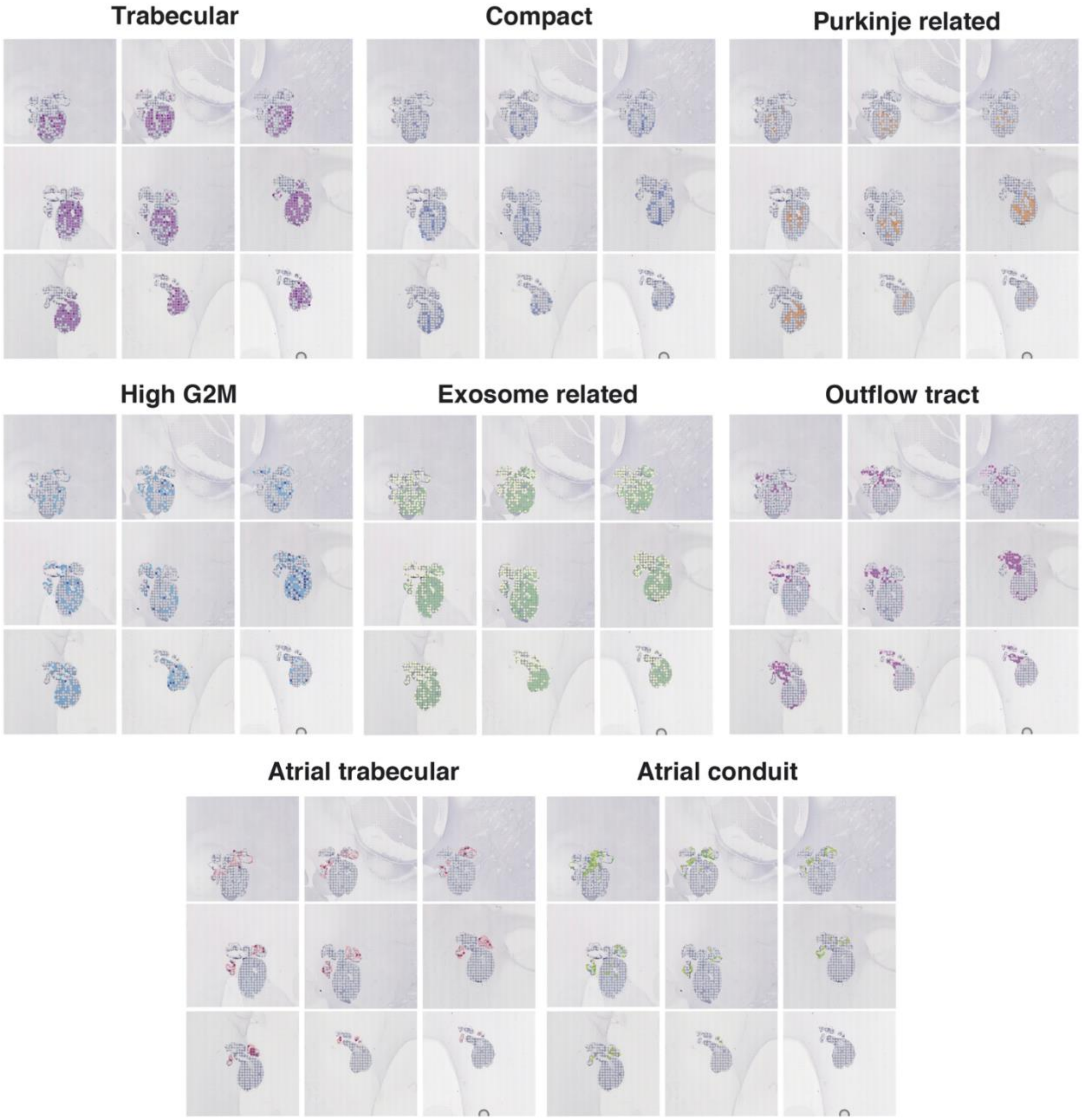

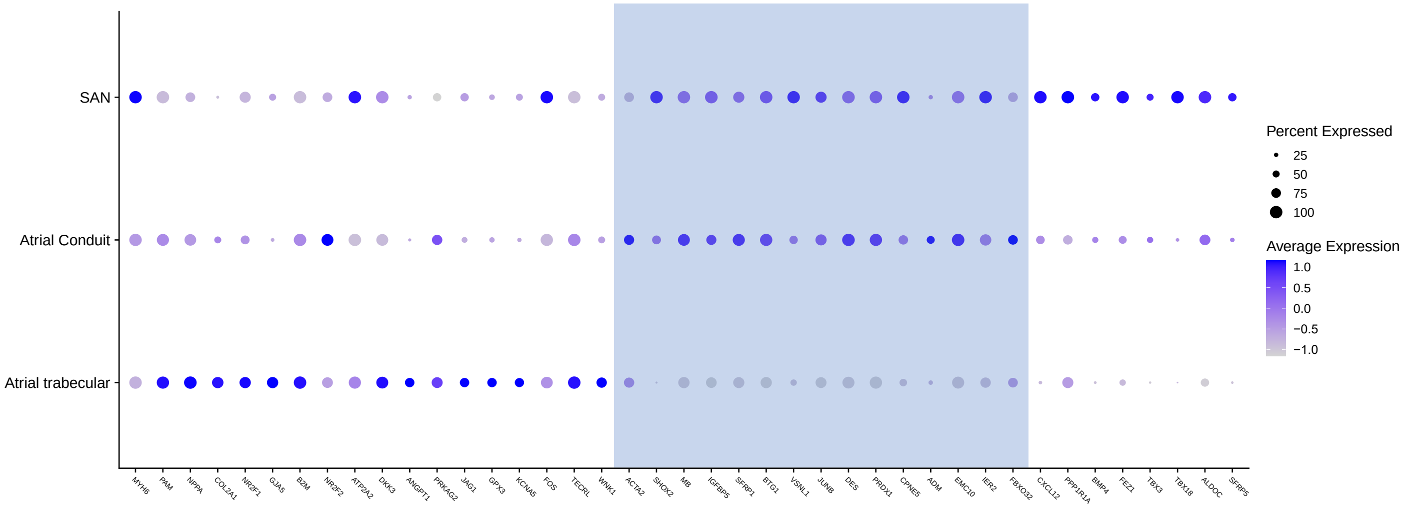

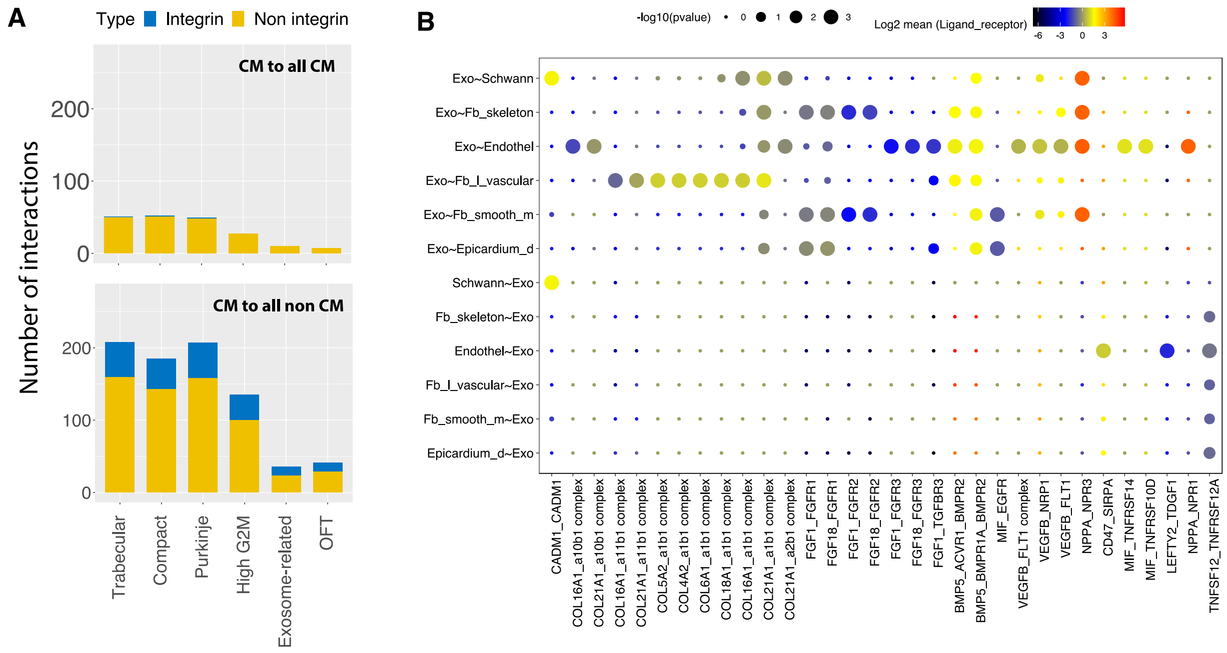

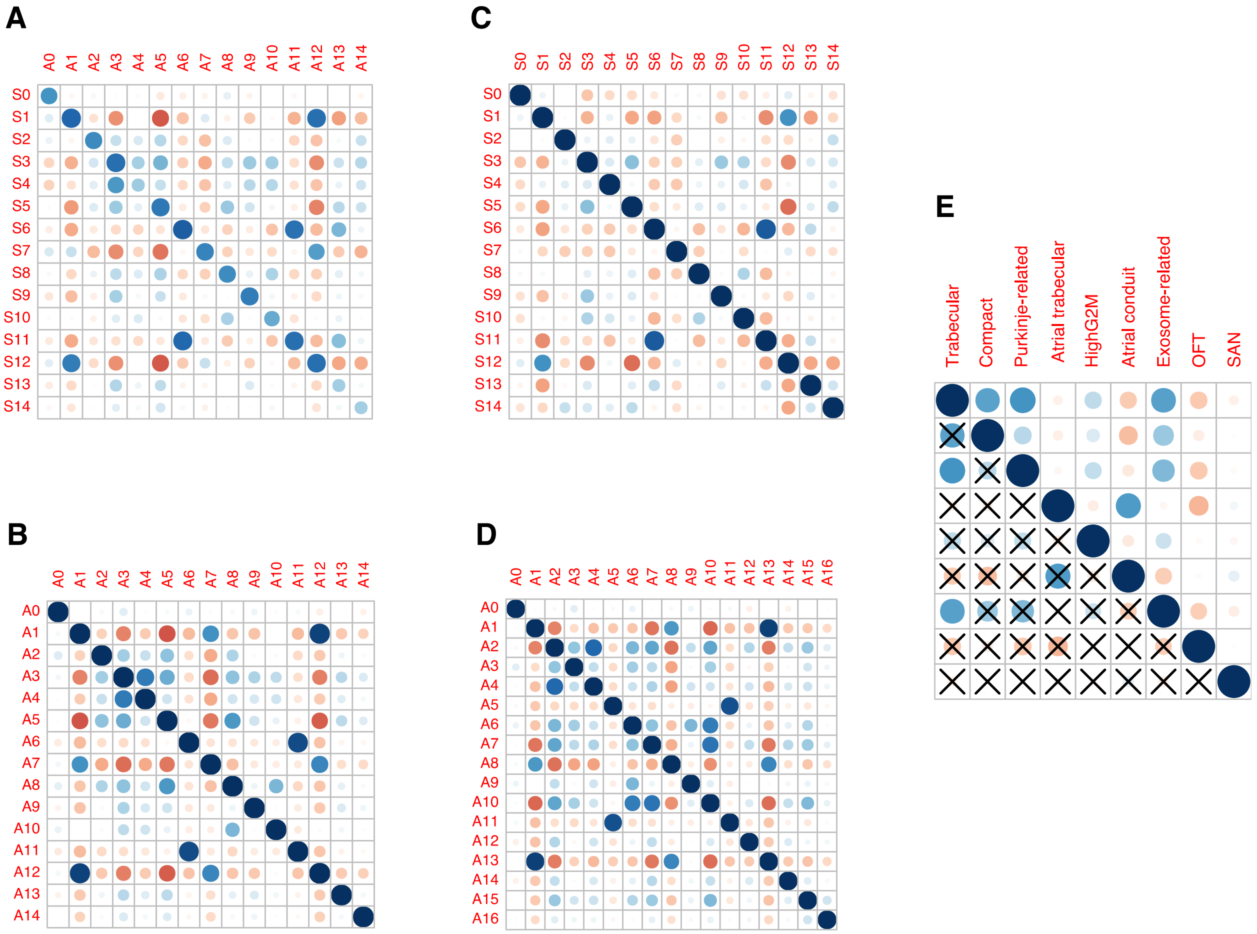
